# Females with epilepsy show abnormal changes to perimenstrual sensory induced long-term potentiation

**DOI:** 10.1101/2025.08.10.669555

**Authors:** Malak Alshakhouri, Cynthia Sharpe, Peter Bergin, Alexander Shaw, Suresh Muthukumaraswamy, Khalid Hamandi, Rachael L. Sumner

## Abstract

**Introduction:** Catamenial epilepsy refers to a cyclical exacerbation of seizure frequency and/or severity linked to stages of the menstrual cycle. To investigate the contribution of underlying excitatory and inhibitory mechanisms of this exacerbation we induced and measured long-term potentiation (LTP). Oestradiol modulates primarily NMDA receptor driven excitation and would enhance visual LTP. Allopregnanolone modulates GABA receptor mediated inhibition and would suppress visual LTP.

**Method:** A control cohort (n = 25) and a cohort with epilepsy (n = 20) were recruited. Participants took part in a visual LTP task while EEG was recorded. Three study visits were timed to capture the mid-follicular (days +5 to +8), mid-luteal (days -9 to -5) phases, as well as the hormone withdrawal in the perimenstrual phase (days -3 to +2). Blood samples confirmed session timing. Generative computational modelling of thalamocortical changes in the EEG was paired with standard evoked potential analysis.

**Result:** 17 (age 26 ± 9.40) participants with epilepsy, and a control cohort of 25 (age 30 ± 7.06) without epilepsy completed the study. The P2 visually evoked potential (a positive-going component of the visual event related potential waveform) was significantly more enhanced by LTP induction in the mid-follicular phase than the mid-luteal or perimenstrual phases in the control cohort (*F*(2, 144) =11.54, p = 0.004 corrected). In contrast, in the epilepsy cohort the P2 was significantly more potentiated in the perimenstrual phase than the mid-follicular or mid-luteal phases (*F*(2,108) = 12.21, p = 0.005 corrected). The effect of cycle on visual LTP was robust, with only 3/17 participants with epilepsy showing highest P2 modulation in the mid-follicular phase (which is the pattern the healthy cohort showed).

The computational modelling showed that in epilepsy, the perimenstrual phase was associated with a decrease in superficial pyramidal grain control as well as decreased LTP driven modulation of feed-forward superficial connections to layer 5 and layer 6 to thalamus.

**Discussion:** Our finding of enhanced visual LTP in the luteal phase and perimenstrual phase of females with epilepsy indicates a breakthrough of glutamatergic excitation. Given the relationship between oestrogen and n-methyl-D-aspartate (NMDA) receptor dependent LTP this is plausibly NMDA receptor driven. Several thalamocortical model parameter changes observed warrant further investigation in studies stratifying participants by catamenial epilepsy type. Overall, while research into allopregnanolone and its inhibitory effects dominate perimenstrual catamenial epilepsy research, this study justifies consideration of the role of seizure exacerbating oestradiol.

## Introduction

Catamenial epilepsy refers to a menstrual cycle caused exacerbation of seizures, typically presenting as a worsening in seizure frequency and/or severity.^1,2^ Three main types of catamenial epilepsy have been described, C1, C2 and C3, according to key stages of the menstrual cycle – menstruation, the follicular phase, ovulation, and the luteal phase.^3^ Type C1 is the most common and the seizure exacerbation occurs perimenstrually. Type C2 refers to a periovulatory exacerbation. Type C3 refers to an exacerbation in the presence of inadequate levels of progesterone in the luteal phase.^3^ C2 and C3 can be attributed to the relatively high oestrogen levels in the perimenstrual and periovulatory phases given that when considering the cortex globally oestradiol lowers seizure threshold.^4^ However, the mechanism behind C1 type is less clear. It is often attributed to the rapid perimenstrual withdrawal of the gammaamino-butyric acid receptor (GABAAR) enhancing neurosteroid allopregnanolone.^5^ Allopregnanolone ordinarily protects against seizures in the luteal phase^4^. This perimenstrual neurosteroidal withdrawal hypothesis^5^ suggests that the brain’s functional response to hormone changes, rather than their absolute concentrations may be key to understanding altered seizure thresholds.

All types of catamenial epilepsy are considered to be due to a global shift in a person’s seizure threshold. Catamenial epilepsy is not a separate epilepsy syndrome rather it manifests as a menstrual-cycle-related exacerbation regardless of the type of epilepsy a patient may have^2^. While C3 catamenial epilepsy has a clear individual-level pathogenesis (inadequate mid-luteal progesterone rise^2^), it is unclear why all females with epilepsy are not equally affected by hormonal shifts over the menstrual cycle in C1 and C2. It is not clear whether the seizure threshold is being changed by an increase in oestrogen driven glutamatergic excitation, or a decrease or dysfunction in allopregnanolone derived GABA-ergic inhibition, or potentially by some other mediator. Animal studies have provided evidence that allopregnanolone withdrawal or inadequate allopregnanolone driven GABAergic inhibition may exacerbate seizures,^5^ and this has led to clinical trials of progesterone^3,6^ (as a prodrug to allopregnanolone) as well as proposals to test the usefulness of allopregnanolone analogue drugs like ganaxolone.^7,8^ The fact that progesterone was found to have only limited usefulness in catamenial epilepsy^3^ and the lack of any effective treatment for catamenial exacerbation of epilepsy^9^ supports the importance of further mechanism-focussed research.^2^

Finding the mechanism underlying catamenial seizure exacerbations may be key to developing better-targeted therapeutic strategies.

Methods for non-invasively detecting changes in excitation and inhibition over the menstrual cycle in humans include transcranial magnetic stimulation (TMS) and electroencephalography (EEG) recordings. Using TMS with paired-pulse inhibition (PPI), studies have shown that females with epilepsy show abnormal PPI changes over the menstrual cycle.^10–13^ In fact, they exhibit the opposite to a healthy control group who demonstrate relatively greater excitation in the midluteal phase compared to the follicular phase, whereas as the control group showed the in the follicular compared to midluteal phase.^10–13^ These findings were not related to different blood-hormone concentrations, suggesting that it is the brain’s response to these sex hormones rather than the hormone level per se. However, the non-specificity of the TMS PPI approach limits inference as to what the underlying mechanisms may be.

Studying LTP provides an opportunity to investigate a specific and ubiquitous excitatory function in the brain relevant to both seizures and the menstrual cycle.^14,15^ Broadly, LTP refers to a strengthening of synaptic connectivity following co-activation of neurons in a network. In vivo induction of LTP typically necessitates that a recording electrode and a stimulating electrode are implanted into a population of cells, and their afferent pathway respectively. The stimulating electrode is used to deliver a presynaptic burst of high frequency electrical stimulation. A subsequent and sustained enhancement of the postsynaptic response (a population excitatory post-synaptic potential) indicates that LTP has been induced in the affected synapse.^16,17^ Visual LTP can be induced using repetitive visual stimulation, resulting in changes in the visually evoked event related potential (ERP) recorded non-invasively at the scalp with EEG.^18^ Modulation of the negative component labelled N1 often occurs in the first minute post-induction and likely reflects a combination of short-term potentiation and early LTP.^18^ The P2 is a positive component of the visually evoked ERP that instead is not potentiated until 30 minutes or more post-induction and likely reflects more sustained early phase LTP changes.^18^

A wealth of *in vitro* and *in vivo* rodent literature suggests that oestrogen can facilitate NMDAR-dependent LTP by enhancing the transmission of NMDARs (likely via the upregulation or receptors) and promoting the formation of synaptic dendrites.^19–22^ Oestrogen can only enhance LTP when it simultaneously enhances dendritic spine density and NMDAR transmission relative to alpha-amino-3-hydroxy-5-methyl-4-isoxazolepropionic acid receptor (AMPAR) transmission^23^. Oestrogen related changes in GABAAR mediated inhibition or GABA release have been shown not to contribute to changes to LTP.

Allopregnanolone, an endogenous neurosteroid, acts as a positive allosteric modulator of GABAARs and would be expected to supress LTP, as GABAAR activity suppresses or prevents LTP induction in neocortex.^24^ In rodents, periods of elevated δ-containing GABAAR expression (the receptor allopregnanolone binds to) were found to be associated with impaired LTP over the oestrus cycle.^25^ Thus studying LTP provides an opportunity to investigate a specific and ubiquitous excitatory function in the brain relevant to both seizures and the menstrual cycle. How LTP changes over the menstrual cycle may reveal relative changes to AMPAR and NMDARs, compared to GABAAR changes and their functional consequences.

Visually induced LTP paradigms have been used to evaluate the effects of female sex-steroids on synaptic plasticity in the context of the naturally occurring menstrual cycle^26^ and oral contraceptive use.^27^ These studies were in a healthy cohort and the perimenstrual phase has never been separated out and tested. There are no reports of the visual LTP paradigm being used in people with epilepsy. The main aim of this study was apply a visual LTP paradigm during three phase of the menstrual cycle – perimenstrual, mid follicular and late-luteal – with a combination of standard ERP and LTP analysis, and a generative computational model of visual cortex to infer the specific glutamatergic and GABAergic mechanisms underlying changes seen across menstrual cycle phases^27,28^ in females with and without epilepsy.

## Methods

### Ethics Approval

The study was approved by the Health and Disability Ethics Committee (Reference number: 21/CEN/201). Participants over the age of 16 provided informed consent, while participants below the age of 16 provided assent along with parent/guardian consent.

### Study Overview

The study was a counterbalanced repeated measure design consisting of three sessions timed to specific points of the menstrual cycle. All sessions were counterbalanced to control for session order effect. A control without epilepsy (n = 25) and an epilepsy (n = 20) cohort were recruited. A CONSORT diagram of recruitment is in Supplementary Material Fig. S1.

Epilepsy participants self-referred and/or were recruited through neurologist and epilepsy specialist nurse referrals. Advertisements were placed in hospitals, GP clinics, university noticeboards, and online.

All participants in the epilepsy cohort were required to keep a seizure diary for at least three months. Participants were asked to report any seizure-like experiences, including ‘auras’ (focal aware seizures). The minimum required information was the date each seizure occurred. Participants are classified by catamenial epilepsy severity as described in^2^ (see Table 3).

**Table 1.** Participants Demographics.

| Characteristic | Controls (*n* = 25) | Epilepsy (*n* = 20) |
| --- | --- | --- |
| Age (years) |  |  |
| Mean ± SD | 30 ± 7.06 | 26 ± 9.40 |
| 14–17 | - | 4 (20.0%) |
| 18–24 | 5 (20%) | 6 (30.0%) |
| 25–34 | 11 (44%) | 5 (25.0%) |
| 35–44 | 6 (24.0%) | 5 (25.0%) |
| 45 and over | - | - |
| Ethnicity, *n* (%) |  |  |
| European | 9 (36.0%) | 15 (75.0%) |
| Māori | 1 (4.0%) | 1 (5.0%) |
| Chinese | 5 (20.0%) | - |
| Indian | 3 (12.0%) | 3 (15.0%) |
| Southeast Asian | 2 (8.0%) | - |
| Middle Eastern | 1 (4.0%) | - |
| Latin American | 1 (4.0%) | 2 (10.0%) |
| African | 1 (4.0%) | - |
| Other | 2 (8.0%) | 1 (5.0%) |

*Note. Total Ethnicity percentages exceed 100% due to multiple ethnicities reported by a single participant. Percentages are based on group totals.*

**Table 2.** Clinical Characteristics of Patients with Epilepsy (*n* = 20)

| **Characteristic** | **$n$ (%)** | **Mean (range)** |
| --- | --- | --- |
| **Age of Onset** |  | 11 (1–31 years) |
| **Epilepsy Duration** |  | 15 (a month – 38 years) |
| **Treatment Type** |  |  |
| Monotherapy | 9 (45.0%) |  |
| Polytherapy | 9 (45.0%) |  |
| None | 2 (10%) |  |
| **Current Medication usage** |  |  |
| Levetiracetam | 8 (38.1%) |  |
| Lamotrigine | 10 (47.6%) |  |
| Clobazam | 4 (19.0%) |  |
| Clonazepam | 1 (4.8%) |  |
| Lacosamide | 3 (14.3%) |  |
| Carbamazepine | 2 (9.5%) |  |
| Sodium Valproate | 1 (4.8%) |  |

*Note. Total Medication percentages exceed 100% as some participants use more than one medication. Percentages are based on the total number of patients.*

**Table 3.**
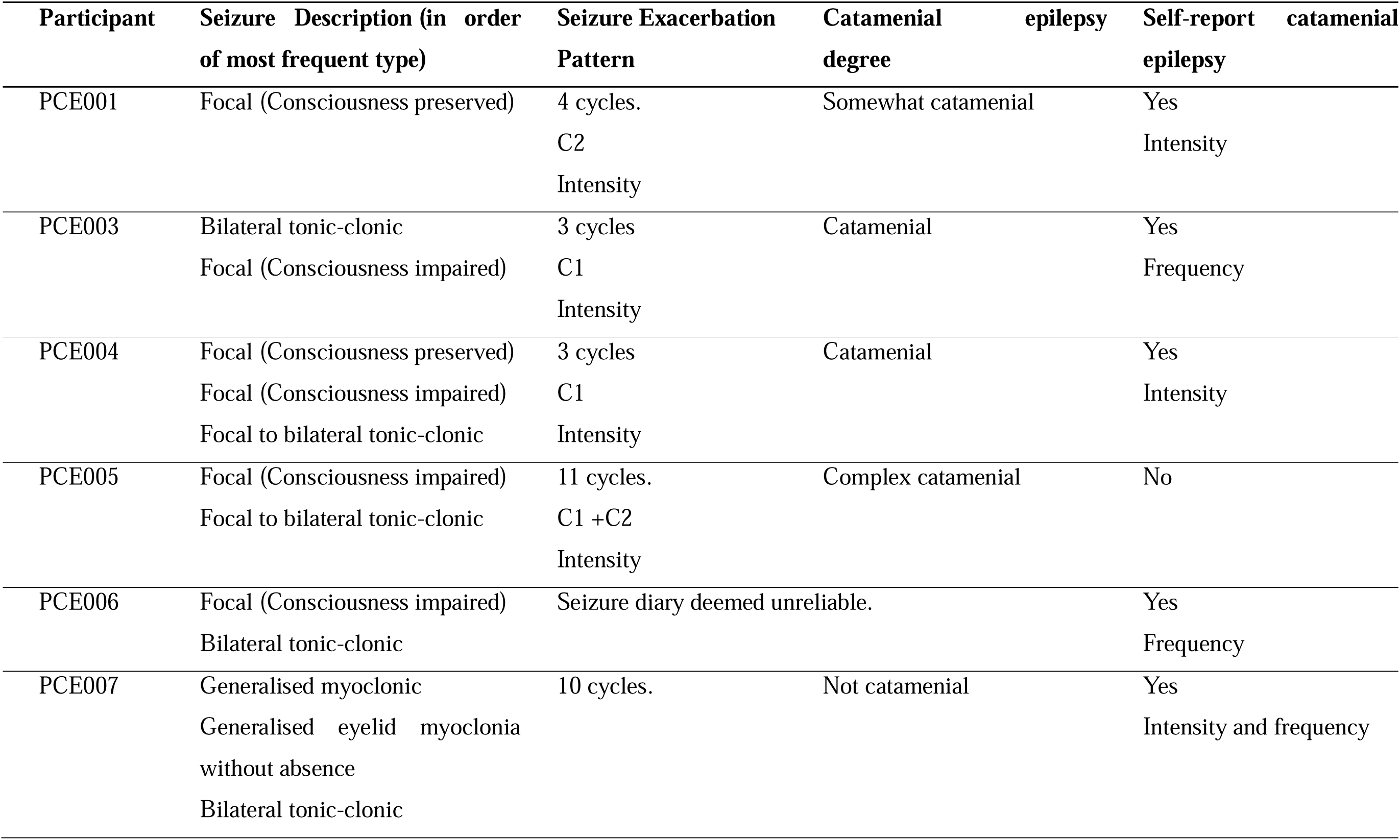
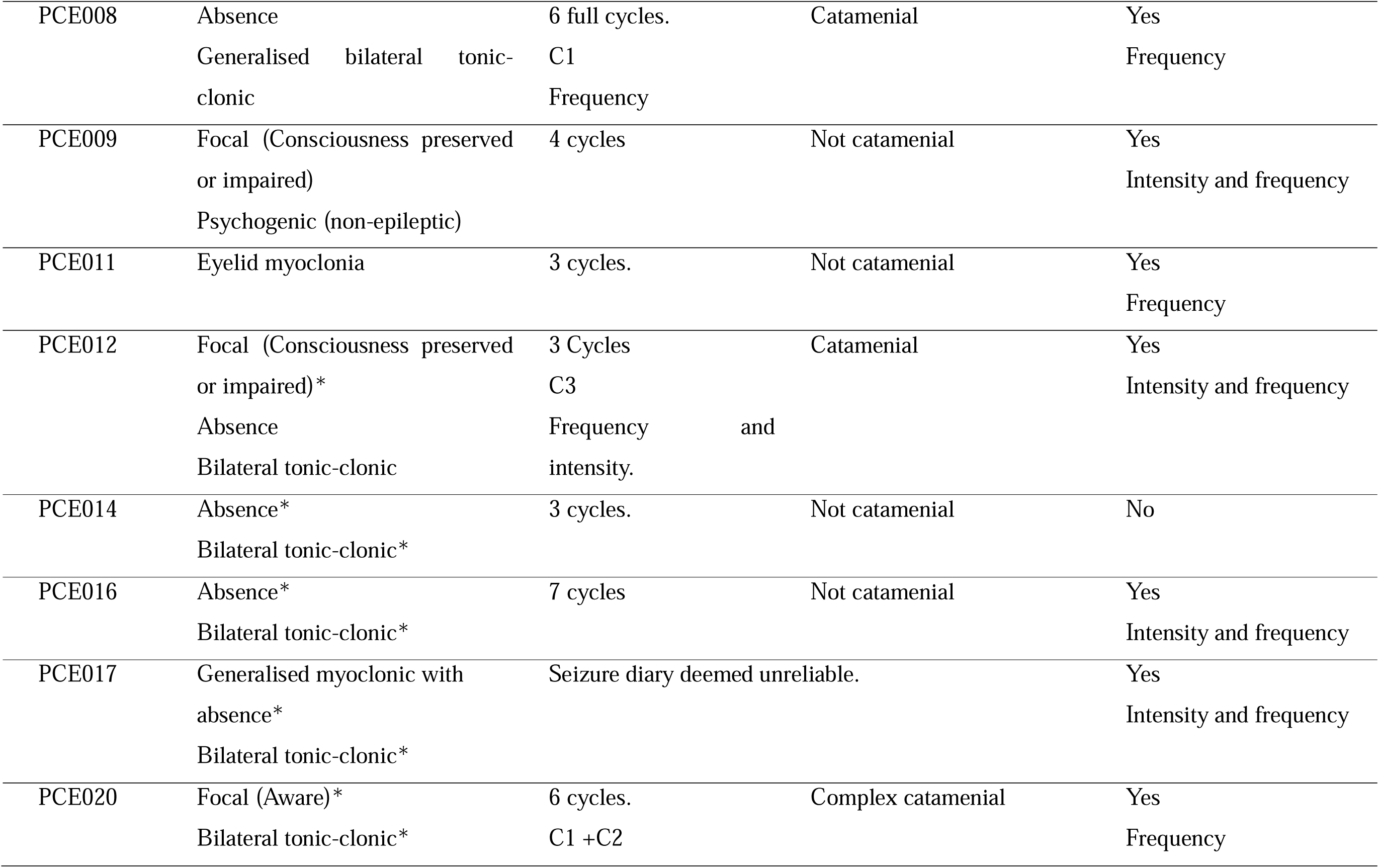
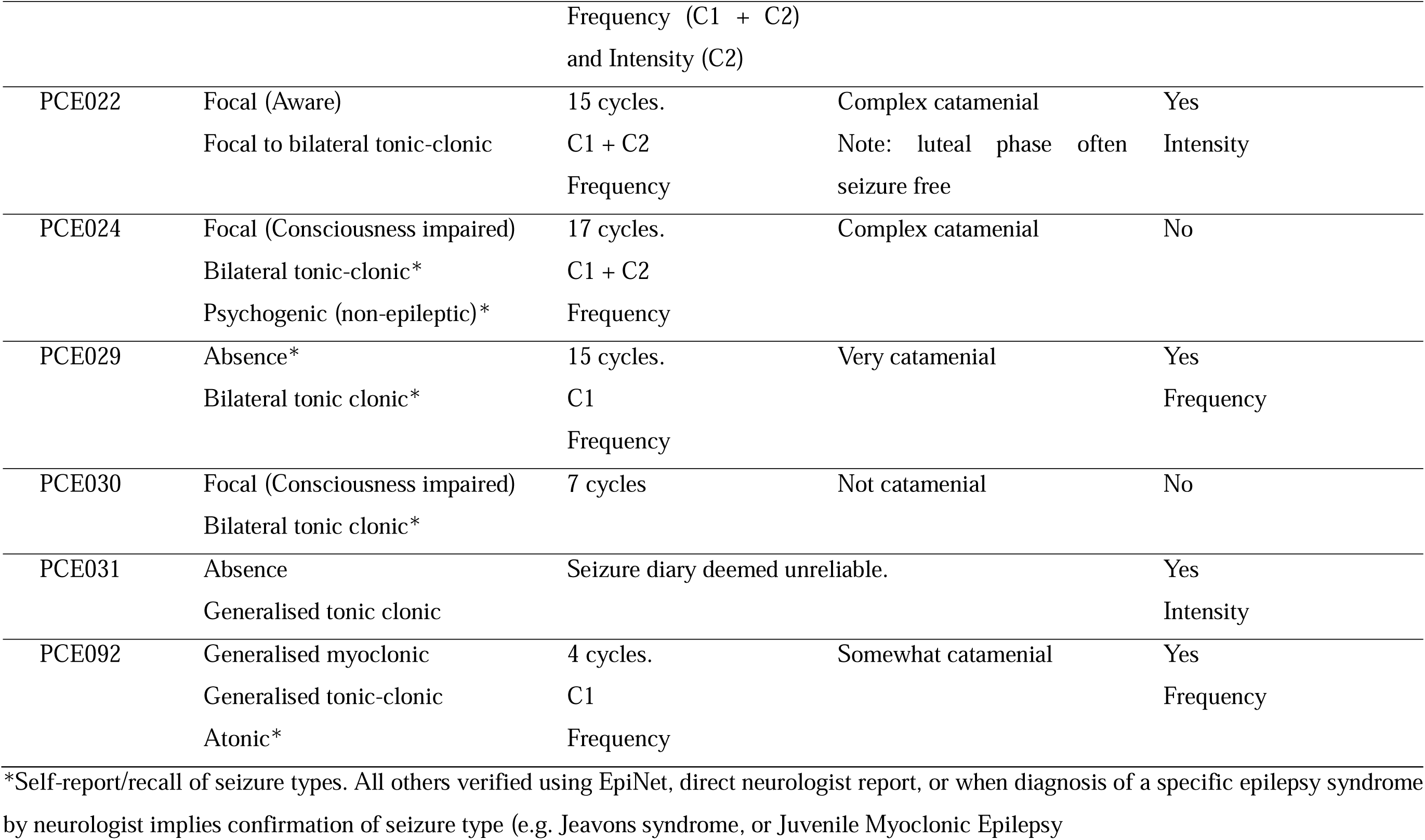
Seizure Characteristics of epilepsy patients.

### Inclusion/Exclusion Criteria

All participants were females aged 14-45 years inclusive, who were willing and able to comply with the study requirements. All participants were required to be stable on any neuroactive medications for at least two months and to have reached a steady state following any dose changes before the first visit (i.e. five half-lives of taking a new dose). They were also required not to change the dose for subsequent visits. Participants with epilepsy must have had their diagnosis confirmed or provided by a neurologist and could be of any type provided they were able to complete the study demands of the EEG and tasks. Comprehensive exclusion criteria can be found in the Supplementary Material.

### EEG Session Timing

The three EEG study sessions were timed to the perimenstrual, midfollicular and mid-luteal phases of the menstrual cycle, as descibed in Box 1. The perimenstrual phase was defined as a 5-day period surrounding the onset of menstrual flow to capture the effects of allopregnanolone withdrawal (days -3 to +2). The mid-follicular phase captures the troughs of progesterone and oestradiol across a 4-day period (days 5 to 8); days 3 and 4 were avoided to provide a clear separation between the perimenstrual and midfollicular sessions. The mid-luteal phase was defined as a 5-day period starting 4 days after the day of ovulation to capture the peaks of progesterone and oestradiol (cycle day -9 to -5). Each session took around 2 hours and started after 12 pm to control for diurnal fluctuations of progesterone and oestradiol.^29^

All participants were required to track their cycle length for at least three cycles prior to the first session to establish the average cycle length for each participant. Several participants, especially in the epilepsy cohort, had considerably short, long, or irregular cycle lengths. To account for cycle length variations and irregularities among those participants, the mid-luteal session was timed 4-7 days after a positive ovulation test. Ovulation testing was completed using digital Clearblue® urine ovulation test. Timing was confirmed on the session day with progesterone level of > 6 nmol/L.

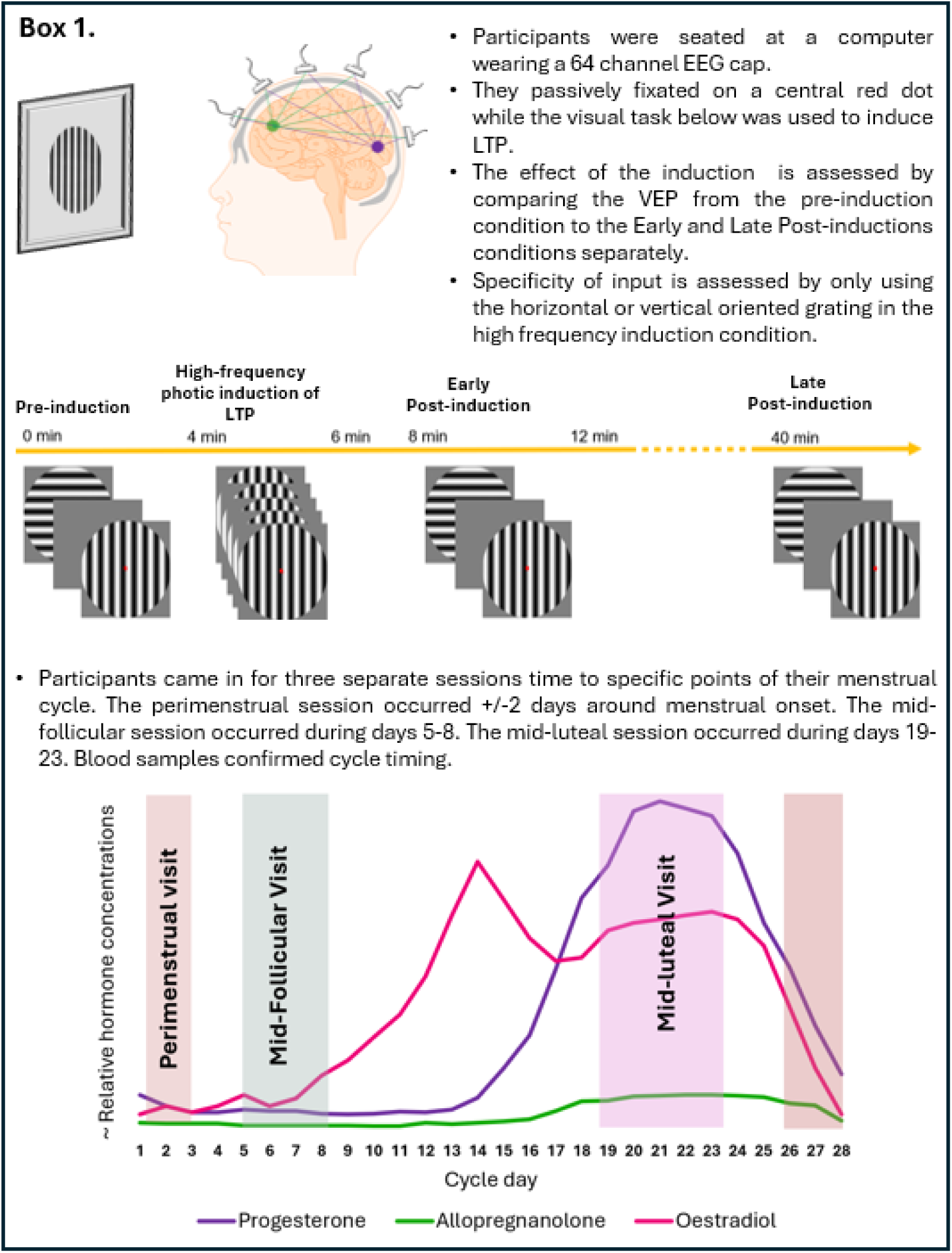

### Demographics and clinical characteristics

The clinical characteristics of the epilepsy cohort were collected in the patients’ EpiNet records (www.epinet.co.nz) and 2. Participants’ study-tracked^2^ as well as self-reporting of catamenial epilepsy at the time of screening (i.e. whether they have noticed a change in seizure frequency and/or intensity during specific phases of their cycle) are provided in Table 3.

### Blood samples

Blood samples were collected successfully from all 25 control participants and 19 epilepsy participants. All blood samples showed progesterone and oestradiol levels that were within the expected range on the study day for the mid-follicular, mid-luteal and perimenstrual cycle phases, confirming correct session timing. One participant confirmed ovulation using urine testing only as they were unable to give a blood sample.

Three separate 2×3 repeated measure ANOVAs ([Group: control vs epilepsy] x [Session: midfollicular vs mid-luteal vs perimenstrual]) have been conducted to test for group differences in each of the measured hormones (progesterone, and oestradiol) during each study session. The results revealed no significant group differences in the measured hormones (all *p*s > .05). See Fig. 1.

**Figure 1.**
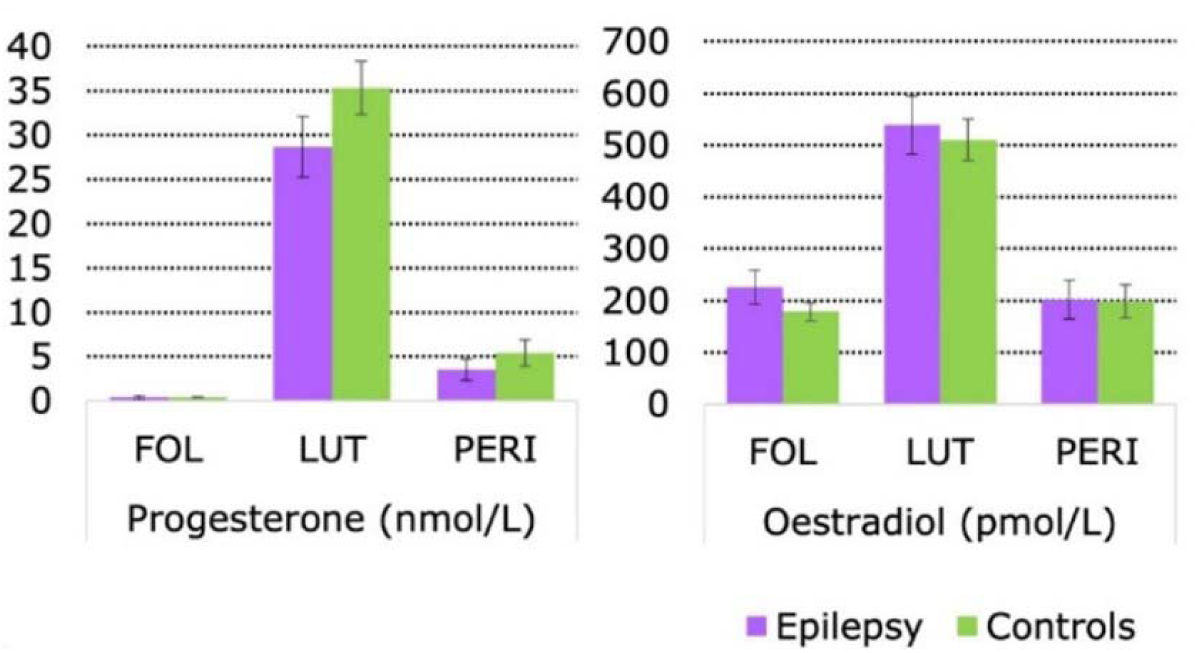
Blood hormone concentrations. Plasma levels of progesterone, and oestradiol per session for Epilepsy (in purple) and controls (in green). Abbreviations: FOL = mid-follicular, LUT = mid-luteal, PERI = perimenstrual.

### EEG Protocol

EEG was recorded using a 64-channel actiCAP with Ag/AgCl active shielded electrodes and Brain Products amplifiers. Participants were comfortably seated in front of a computer screen. To visually induce LTP they were shown intermittent horizontal and vertical circular sine gratings across four conditions (Box 1). Participants were instructed to passively fixate on a red dot presented in the centre of the screen throughout the task. In the pre-induction condition stimuli were presented 120 times each in random order at low frequency (1 Hz), for a total duration of 4 minutes. The purpose of the pre-induction condition was to establish a baseline ERP amplitude for subsequent comparison with post-induction conditions. The second condition was a 2-minute photic induction of LTP involving 1000 presentations of either the horizontal or vertical orientation at a higher frequency (9 Hz) to induce LTP. The orientation of the stimulus used in the photic induction was counterbalanced between participants to test for input specificity. Post-induction there are two repetitions of the baseline condition to assess ERP modulation early (2-minutes) and late (40-minutes) postinduction.

### EEG Analysis

25 control cohort participants and 17 epilepsy cohort participants were included in this analysis. Two with epilepsy were excluded due poor data quality, one was excluded as ovulation was not evident during the study period. EEG preprocessing was carried out using Fieldtrip toolbox.^30^ See the Supplementary Material for full details. Difference waves are calculated for all statistical analysis where the pre-induction baseline average was subtracted from the early post-induction and late post-induction for all participants. For all analyses, main effects were considered significant at p < 0.05 family-wise error corrected (FWE-c). Simple effects tests were conducted as appropriate. Any posthoc contrasts are FWE-c. Where multiple significant ERP peaks occurred for the same component, only the most significant peak within a cluster is reported.

For each cohort, the initial ERP parameter finding analysis was a 3×2×2 repeated-measure analysis of variance (ANOVA) run across a 0-250 ms time window ([Phase: mid-follicular vs luteal vs perimenstrual] x [Time: Early Post-induction vs Late Post-induction difference waves] x [Specificity: horizontal or vertical stimulus orientation for induction vs not used]). The purpose of this analysis was to confirm that high frequency photic induction of LTP had induced typical modulation of the visually evoked potential^18^ and to identify parameters for follow-up analyses. Post hoc t-contrasts were used to assess the direction of change for each component. All other interactions and simple effects were considered irrelevant for the purpose of this parameter-finding analysis.

### Computational modelling

EEG source analysis located the peak voxel in visual cortex representing the effect of visually inducing LTP on the EEG and is explained in detail in the Supplementary Material. The virtual local field potential (LFP) was extracted as a 5 mm sphere and the generative Thalamocortical Model (TCM) model was fit to the LFP timeseries [0-250 ms] using Dynamic Causal Modelling (DCM).^31^ The model architecture is fully depicted in the Supplementary Fig. S2.

The model inversion protocol incorporated a parametrised general linear model on synaptic parameters capturing slow between-stage dynamics. The effect of LTP was modelled as a linear change from baseline to the Late condition [-1 0 1] as well as a non-linear change from baseline to the Early condition [-1 1 0]. The linear contrast reflects early phase LTP and the time course for P2 potentiation, while the non-linear contrast models general excitability or short-term potentiation and the time course for N1 potentiation. These two contrasts were then combined in [-1 1 0; -1 0 1] contrast to allow for both non-linear and linear contributions to describe the condition-specific effects, representing the model that best fit in a methodological investigation to describe visually induced LTP.^31^

Estimation of the parameters of primary visual cortex microcircuitry contributing to the effect of menstrual cycle on LTP was conducted using Parametric Empirical Bayes (PEB).^32^ A threshold of ‘very strong evidence was applied to the remaining parameters by only reporting parameters change with posterior probability (Pp) > 0.99. For more detail see Supplementary Material.

### Study One, Two and Three: cohort and catamenial classification

The cohort of females without epilepsy is reported as Study One, and the entire cohort with epilepsy forms Study Two. Also in Study Two, to assess whether visual LTP was different between the cohort with and without epilepsy a 3×2 repeated measure ANOVA ([Phase: midfollicular vs mid-luteal vs perimenstrual] x [Group: control vs epilepsy]) was performed. Study Three added the effect of catamenial epilepsy to the between group differences. Given there are only 17 participants with epilepsy and useable EEG and seizure tracking data this was a qualitative assessment only.

## Results

### Study One: Females without epilepsy

#### ERP results

The initial parameter finding analysis specifically assessed main effects of time on negative-going components in the early post-induction condition and positive-going components in the late post-induction condition, as well as main effects of specificity in both conditions targeting the most common findings of ERP modulation in previous studies (reviewed in^18^). The negative early potentiation contrast revealed no significant effects of time, indicating that tetanus induced no significant changes in the negative-going components of the ERP. However, there was a significant interaction between specificity and time in the early post-induction condition, that included a centro-parietal peak at 132ms (*F*(1,288) = 24.03, *p* = 0.001 FWE-c). Follow up testing of this two-way interaction revealed no significant effects (all *p*s > .05), indicating that there was no clear difference in enhanced negativity between the induced stimulus orientation and non-induced stimulus orientation that could be described as specificity.

In contrast, the positive late potentiation contrast revealed a significant effect of time that included a centralised cluster extending from 183-244ms and peaking at 183ms (*t*(288) = 7.61, *p* = 3.80 ×10^−10^ FWE-c). The distribution and timing of this cluster appear to be consistent with the central distribution of the P2 component (Fig. 2A). No interaction between post-induction time was found in the late post-induction block, indicating that P2 potentiation did not demonstrate specificity to tetanising stimulus orientation.

**Figure 2.**
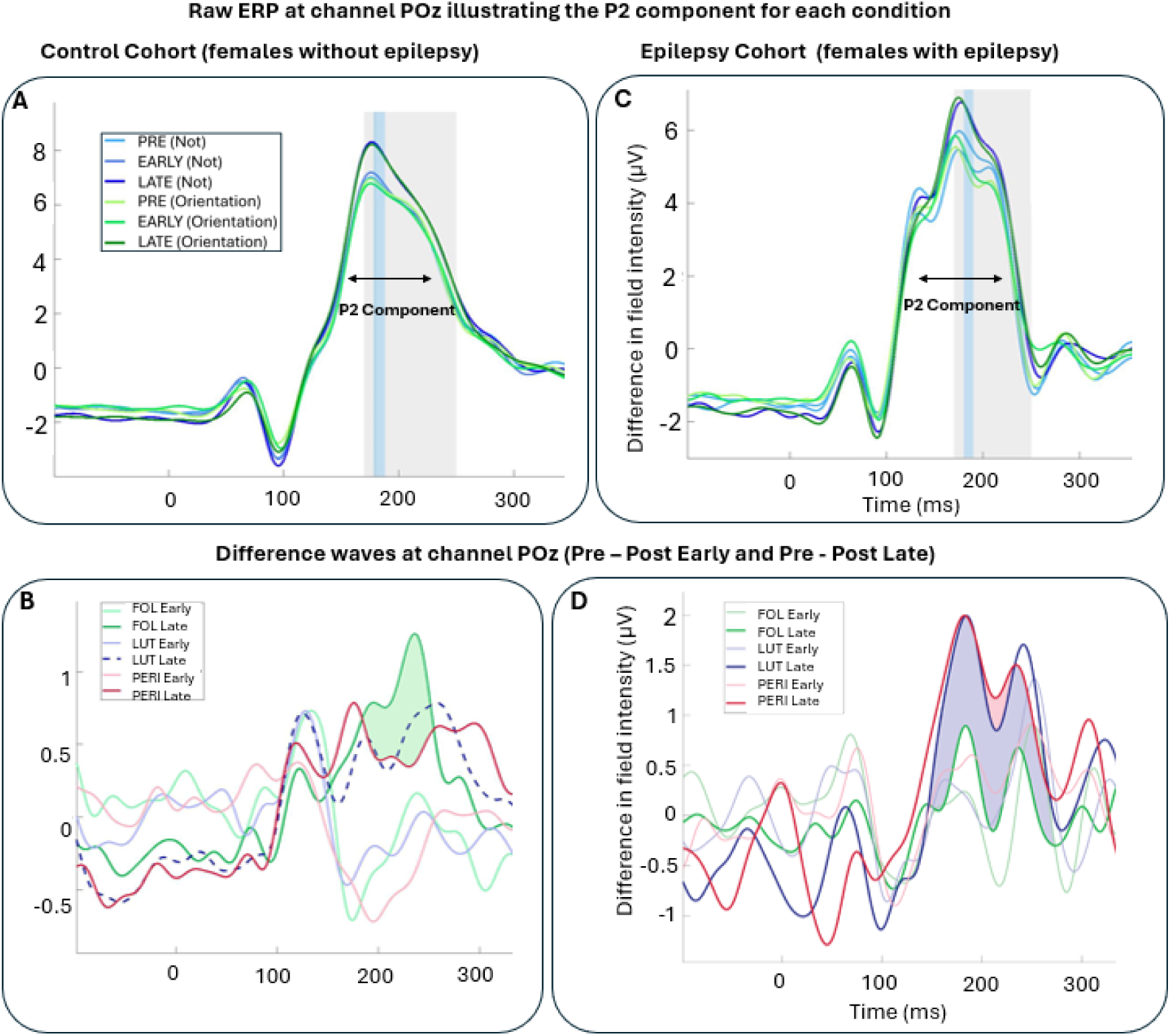
Event related potentials and difference waves showing visually induced LTP effects on the P2 evoked component for both cohorts. Raw grand average ERP waveform plotted for all participants show the ERP and LTP data look similar in the control and epilepsy cohort. A) shows the control cohort and the B) epilepsy cohort. The timing of the significant peak where the P2 is most enhanced (dark lines higher than light lines) is shaded in blue and the time window of interest for the follow-up analysis is shaded in grey. C) For healthy controls Early and Late time difference waves at channel POz showing a significant interaction between Cycle Phase and Time in the late post-tetanus block, but not the early post-tetanus block. Green shaded area reflects the significant difference between the late midfollicular phase (dark green) and late perimenstrual phase (red). The late mid-luteal phase (dashed dark blue) did not differ significantly different from the other two phases. No significant differences were found between the early time difference waves between any of the phases. D) For the cohort with epilepsy blue shaded area reflects the significant difference between the late mid-luteal phase (dark blue) and late mid-follicular phase (dark green). Pink shaded area shows that the late perimenstrual phase is even more significantly different from the late mid-follicular phase (dark green) than the late mid-luteal phase (dark blue). No significant differences were found between the early time difference waves between any of the phases.

The results of this initial analysis were used to inform the follow-up analyses. Choosing a narrow time window reduces multiple comparisons, thus for the follow up analysis a time window of interest was chosen as 170-250 ms to capture the P2 cluster that has been identified (183-244 ms). The main effect of menstrual cycle on LTP was explored in the late post-induction block where reliable LTP was found. This was run as a 3×2 repeated-measure ANOVA ([Phase: mid-follicular vs luteal vs perimenstrual] x [Specificity: tetanised vs non-tetanised) on the late difference waves. We found a significant interaction between cycle phase and time in the late post-tetanus condition, capturing a left-lateralised increase in ERP modulation at 233 ms (*F*(2, 144) =11.54, *p* = 0.004 FWE-c). This interaction between cycle phase and time indicate that the magnitude of P2 potentiation differed between cycle phases.

Post-hoc testing revealed that this interaction effect is best explained by a significant increase in P2 potentiation during the follicular phase compared to the perimenstrual phase (*t*(144) = 4.72, p = 0.001 FWE-c) (Fig. 2B).There was no significant differences in P2 potentiation between the mid-luteal phase and the mid-follicular phase nor the mid-luteal phase and the perimenstrual phase (all *p*s > .05).

#### Computational modelling

The selected peak was extracted from each participant as the local field potential (LFP) at MNI coordinate [-8 -98 -12] in left middle occipital gyrus. The combination model provided an excellent fit (average variance explained > %95.6, SD = 0.083). Presented in Fig. 3, PEB analysis revealed increased visual LTP task driven modulation of the excitatory to inhibitory ss→si connections (Effect size, Ep = 0.079, Pp = 1.00) and inhibitory intrinsic si→si self-gain (Ep = 0.098, Pp = 1.00) in the perimenstrual session relative to the mid-follicular session.

**Figure 3.**
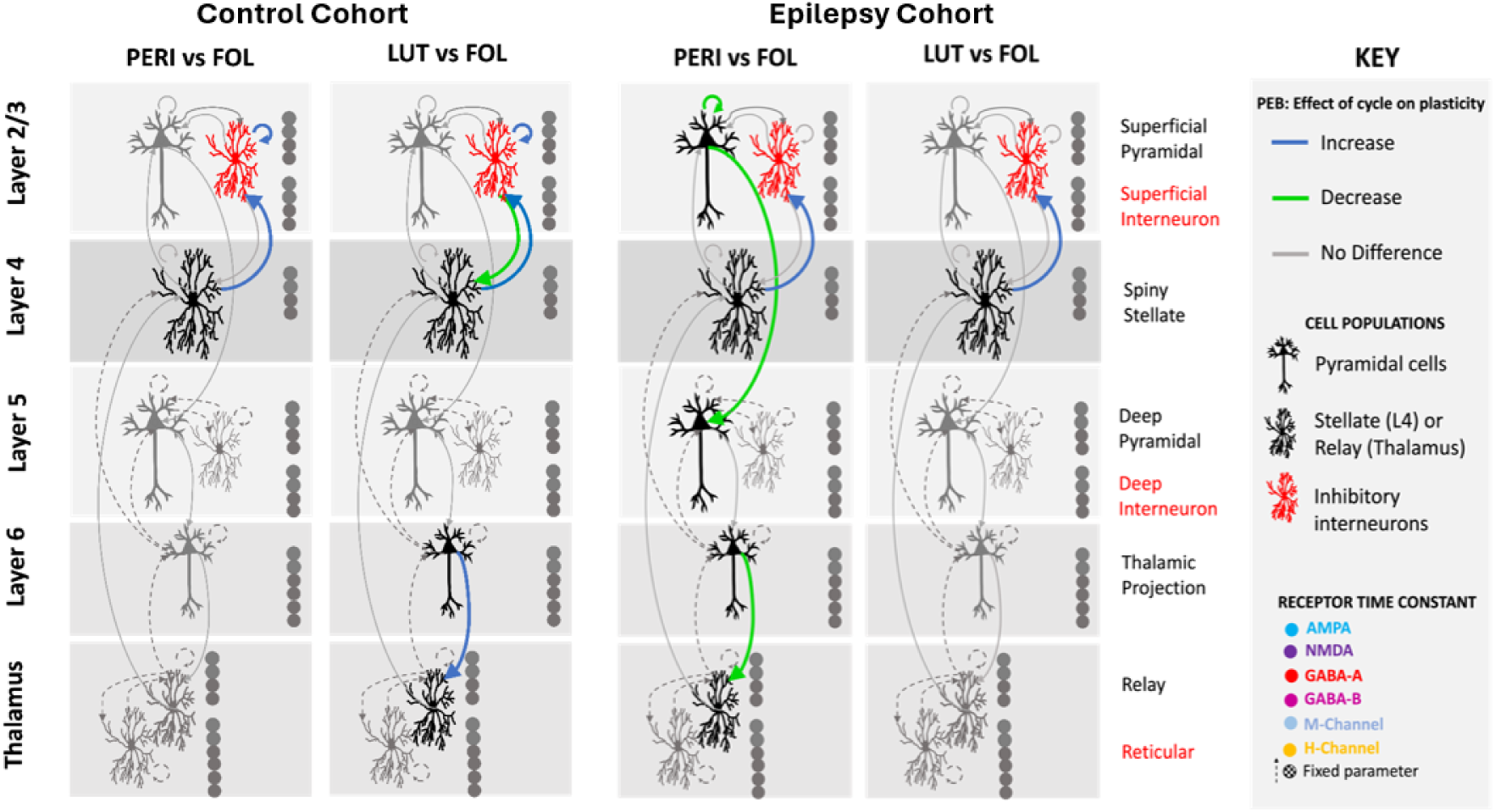
Thalamocortical modelling. Parameters with very strong evidence showing the difference in LTP modulation for the perimenstrual session (PERI) relative to the midfollicular (FOL) and the mid-luteal session (LUT) relative to the mid-follicular session (FOL). Green indicates a parameter was modulated significantly less relative to FOL. Blue indicates a parameter was modulated significantly more relative to FOL.

Additionally, while the ERP analyses revealed no significant differences in the mid-luteal phase compared to the mid-follicular phase, the DCM modelling showed a combination of significantly increased inhibitory parameter si→si (Ep = 0.097, Pp = 1.00), increased excitatory parameters ss→si (Ep = 0.074, Pp = 1.00) and tp→rl (Ep = 0.065, Pp = 1.00) and reduced inhibitory parameter si→ss (Ep = -0.13, Pp = 1.00) in the mid-luteal phase compared to the mid-follicular phase.

In line with the ERP analyses, the DCM modelling showed no significant differences between the mid-luteal phase and the perimenstrual phase.

### Study Two: Females with epilepsy compared to healthy controls

#### ERP results

Following the same order of analyses as the control cohort, the initial analysis was run on the full-time window (0-250 ms) and was conducted as 3×2×2 repeated measure ANOVA. There was no significant effect of time on the early negative ERP components. Nor there was an effect of specificity in the early post-induction or the late post-induction blocks. However, there was a significant effect of time on the modulation of the late ERP components that included a central cluster starting and peaking at 185ms and extending until 236ms (*t*(216) = 9.39, *p* < 0.01 FWE-c). This cluster appear to represent the modulation of the P2 component (Fig. 2C)

Of particular interest was the significant interaction between time and phase that included a slightly left-lateralised parietal peak at 199ms (*F*(2,216) = 12.81, *p* = 0.005 FWE-c) and a small right-lateralised cluster peaking at 6ms. The peak at 199ms appear to represent an effect of menstrual cycle on late LTP modulation. The 6ms peak is too early to represent visual processing and is disregarded as noise. While this interaction was significant, the planned follow-up repeated measure ANOVA was still conducted on the late post-tetanus block to further investigate this interaction.

The menstrual cycle effect on late potentiation was probed as a 3 x 2 repeated-measure ANOVA again as in Study 1 that included only the late difference waves. The time window of interest was identified as 170-250 ms to capture the P2 cluster identified earlier (185ms-236 ms) and to keep it consistent between cohorts. This analysis revealed a significant cycle effect that captured the same peak at 199ms (*F*(2,108) = 12.21, *p* = 0.005 FWE-c).

Post hoc t-contrasts revealed that this interaction is underlined by significant differences in P2 potentiation during the mid-follicular phase relative to the perimenstrual phase (at 200ms; *F*(1,108) = 4.21, *p* = 0.003 FWE-c) and to the mid-luteal phase (at 198ms; *F*(1,108) = 19.78, *p* = 0.005 FWE-c) (Fig. 2D). The perimenstrual phase was associated with significantly greater P2 potentiation than the mid-follicular phase (*t*(108) = 4.58, *p* = 0.002 FWE-c). Similarly, the mid-luteal phase showed greater P2 potentiation than the mid-follicular phase (*t*(108) = 4.45, *p* = 0.003 FWE-c). There were no significant differences in P2 potentiation between the mid-luteal and perimenstrual phases.

#### Computational modelling

Detailed source results can be found in the Supplementary Material. The most significant peak corresponded to MNI = [24 -98 -12] in the right middle occipital gyrus and was extracted as the virtual LFP. The model provided an excellent fit with average variance explained >%94, SD = 0.084. PEB analysis revealed significant changes in parameters between the mid-follicular phase and the perimenstrual phase, as well as between the mid-follicular phase and the mid-luteal phase. The parameters that survived the threshold for strong evidence in the perimenstrual phase relative to the mid-follicular phase involved decreased modulation of excitatory feedforward sp→dp connections (Ep = -0.089, Pp = 1.00), and tp→rl connections (Ep = -0.061, Pp = 1.00), inhibitory sp→sp self-gain (Ep = - 0.17, Pp = 1.00), as well as increased modulation of excitatory ss→si connections (Ep = 0.066, Pp = 1.00).

Additionally, in the mid-luteal phase compared to the mid-follicular phase there was an increased modulation of excitatory ss→si connections (Ep = 0.072, Pp = 1.00). There were no significant differences in the LTP driven modulation of the underlying circuitry between the mid-luteal and perimenstrual phases. See Fig. 3.

### Study Three: Between group (epilepsy vs without) and within group (seizure exacerbation) driven differences

Catamenial epilepsy classification is presented in Table 3. We followed recommendations published in Alshakhouri, Sharpe, Bergin and Sumner ^2^ for classification. However, in order to be transparent about the complexity of classifying catamenial epilepsy we provided further detail in each case. Participants rated as “Somewhat Catamenial” are : (1) participants in whom 50% or more cycles were catamenial, (2) we tracked for longer than the 3-cycle criteria, and (3) they could have reasonably received a catamenial diagnosis if they had only tracked for 3 cycles per the standard criteria.^2,3^ Complex catamenial means they met the criteria for both C1 + C2 type but did not always show both in the same cycles and this makes them somewhat catamenial for one and/or the other in isolation. Catamenial or very catamenial means they either met the recommended criteria^2,3^ of being catamenial for two thirds of cycles or greatly exceeded it.

#### Between group difference in LTP magnitude

A 3×2 repeated measure ANOVA was performed to assess differences in LTP magnitude between groups. There was no significant effect of group on LTP. This is evident in Fig. 2 A and C where the ERP and modulation look similar overall.

Catamenial epilepsy effects on LTP. In order to visualise the effect of catamenial seizure exacerbation on visual LTP at the individual participant level the change in the amplitude of the P2 peak from pre-induction to late post-induction was computed for each subject per session. The P2 peak was manually identified for each participant to account for inter-individual variability in the timing of the P2 peak. Manual identification involved inspecting each participant’s ERP waveform and topography. The P2 peak was defined as a 50 ms around the maximum peak within the time-window of interest (170-250 ms).

The computed P2 potentiation values are plotted as bar graphs in Fig. 4. For only three participants P2 potentiation is highest in the mid-follicular phase compared to the mid-luteal and perimenstrual phase (as in the control cohort main result). Table 3 compared with Fig. 4 shows that the order of phases with greatest P2 potentiation appears randomly distributed among participants with catamenial exacerbations of different types.

**Figure 4.**
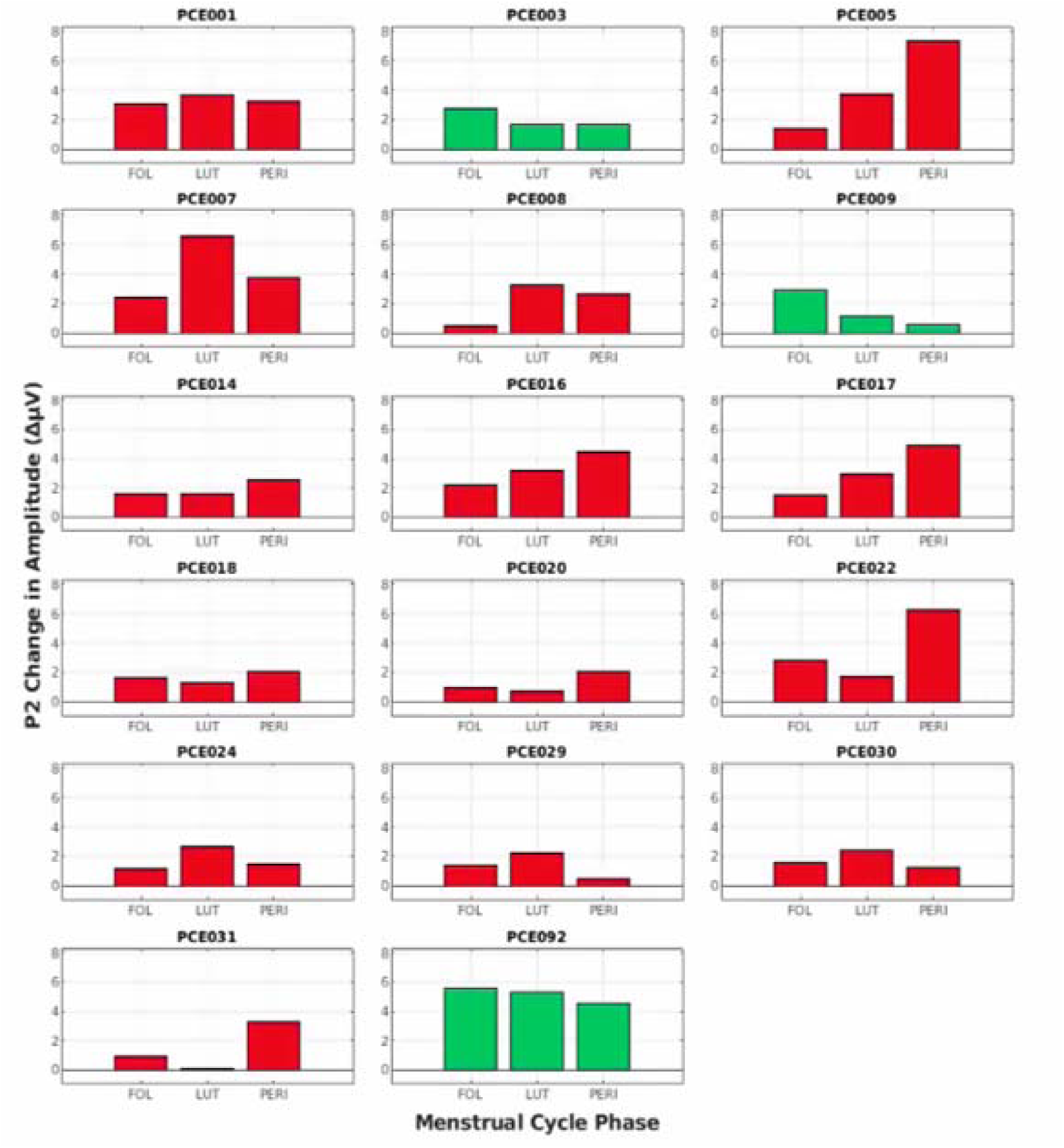
Individual P2 amplitude changes across the menstrual cycle. Individual changes in P2 potentiation from pre-induction of LTP to late post-induction across the menstrual cycle. Red indicates that the magnitude of P2 potentiation is the lowest during the midfollicular phase compared to the perimenstrual and/or mid-luteal phase. Green indicates that the change in P2 peak is the highest during the mid-follicular phase compared to the perimenstrual and/or mid-luteal phases.

## Discussion□

In this study LTP was visually induced and recorded as the consequent enhancement of ERPs. In females with and without epilepsy we recorded EEG during three phases of the menstrual cycle and compared the perimenstrual phase with the mid-follicular and late-luteal phases to capture the effects of neurosteroid withdrawal from female sex steroids. We stratified the epilepsy cohort by catamenial epilepsy type to explore the relationship between changes in visually induced LTP over the menstrual cycle and seizure exacerbation. Study One demonstrated that in female controls without epilepsy visual LTP in the perimenstrual phase is suppressed compared to the mid-follicular phase (induced P2 modulation is less); it appears most like the mid-luteal phase. Study Two investigated changes in visual LTP across the menstrual cycle of females with epilepsy and revealed a trend in the opposite direction to the control cohort, where the perimenstrual phase was associated with greater visual LTP (induced P2 potentiation) than the mid-follicular phase. This key finding is illustrated in Box 2. Study Three investigated the relationship between changes in visual LTP across the menstrual cycle and catamenial epilepsy pattern and found that type of catamenial epilepsy did not specifically related to changes in LTP over the menstrual cycle. The individual level results do however demonstrate how robust the effect of cycle on visual LTP was across individuals with epilepsy.

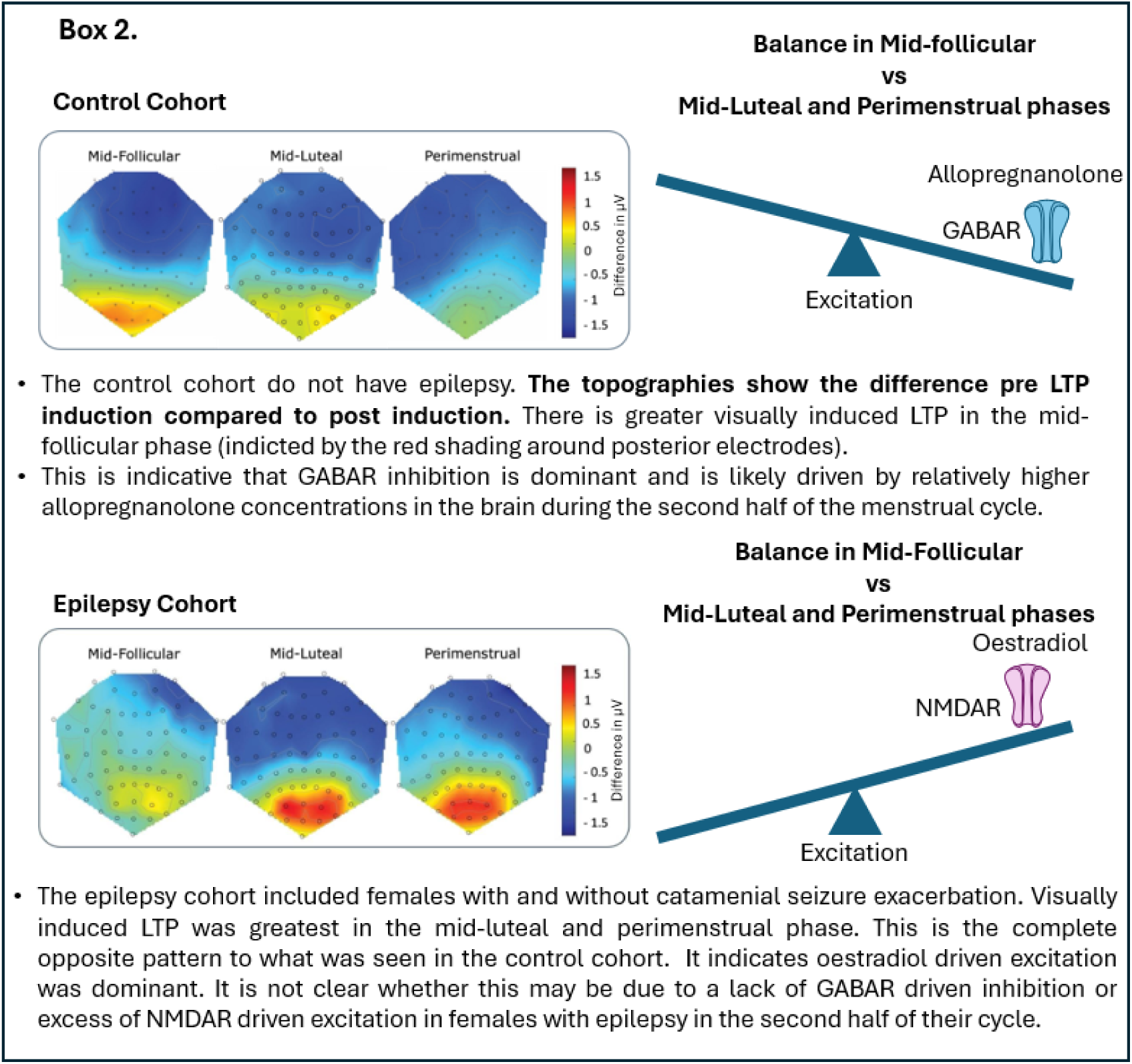

In particular, this study highlights the importance of examining functional changes over the menstrual cycle. Typically, studies time investigations (such as study visits) based on absolute hormone concentration changes. For example, mid-luteal versus mid-follicular versus ovulation. The perimenstrual phase has mean hormone levels as low, or almost as low, as the mid-follicular phase (Fig. 1); yet in the current study the mid-follicular phase and perimenstrual phase showed significant differences in ERP and computation modelling results, while the mid-luteal and the perimenstrual phases were not significantly different in the control cohort. This distinction is particularly important for understanding premenstrual disorders that worsen during the neurosteroid withdrawal, such as C1 catamenial epilepsy. But also menstrual migraine and premenstrual dysphoric disorder.

### ERP findings

Sumner, Spriggs, McMillan, Sundram, Kirk and Muthukumaraswamy ^33^ used the same visual LTP paradigm as in the current study, administered during the early follicular phase (days 1-5) and the mid-luteal phase (days 20-25). They found only a weak effect of the menstrual cycle on visual LTP and speculated the session timing (encompassing the perimenstrual phase across the follicular/luteal phases) may explain the weakness. Therefore, we modified our session timing to provide clear separation from the perimenstrual phase and tested the perimenstrual phase separately. Our research indeed produced a consistent but much more robust effect of the menstrual cycle on visual LTP. Further, it revealed that the menstrual cycle modulation of LTP during the perimenstrual phase (which includes the early follicular phase (days 1-2)) resembled the mid-luteal phase, rather than the mid-follicular phase (days 5-8).

This study demonstrates the importance of the perimenstrual phase in the brain. Further that in humans the effects of the luteal phase are still evident in early follicular phase. This is consistent with the temporal profile of menstrual cycle disorders that tend to resolve within the first few days of menstruation.^34^ The study also demonstrates that visually induced LTP is sensitive to the functional change that is occurring.

Opposite to the control cohort, in females with epilepsy, the mid-follicular phase was associated with the least LTP induced enhancement of the P2 component. The magnitude of P2 potentiation increased significantly in the mid-luteal phase and remained increased in the perimenstrual phase relative to the mid-follicular phase. No significant differences in P2 potentiation were found between the mid-luteal and perimenstrual phases. Inspection of individual LTP patterns revealed that all but three of our epilepsy participants exhibited LTP patterns distinct from that observed in the control cohort. This suggests that LTP magnitude does not systematically vary in accordance with catamenial epilepsy patterns. Instead, our finding of reversed LTP pattern in epilepsy may reflect a broader epilepsy-related neurophysiological trait. This is consistent with what had been found using TMS.^10,12^ Overall this suggests that there is a glutamate mediated and NMDA dependent, abnormal, change in excitation/excitability over the menstrual cycle in females with epilepsy.

Given there was no significant difference in hormone concentrations across the groups and all included participants with epilepsy had ordinary ovulation when tested - our data support that the neurons of females with epilepsy may respond abnormally to typical endocrine events of the mid-luteal phase. Changes in sensitivity to hormones have also been shown in premenstrual dysphoric disorder,^35,36^ and thus could likewise underlie catamenial exacerbation of epilepsy.

The neurosteroid withdrawal hypothesis of catamenial epilepsy posits that the long-term exposure to allopregnanolone of the mid-luteal phase leads to an upregulation of GABAARs that contain the δ subunit.^2^ The withdrawal of allopregnanolone causes further upregulation.^2^ Though never demonstrated in humans, in females without epilepsy this is implied to cause is a net increase in GABAergic inhibition. This is consistent with our finding of decreased LTP in the mid-luteal phase that is also apparent in the perimenstrual phase. Catamenial epilepsy is said to be caused by some dysregulation of GABAAR-δ driven inhibition. Our finding of enhanced visual LTP in the luteal phase and perimenstrual phase of patients with epilepsy supports the theory of dysregulation of GABAAR-δ driven inhibition in all females with epilepsy across the menstrual cycle. However more distinctly, our results strongly indicate there is a breakthrough of glutamatergic excitation. Given the relationship between oestradiol and NMDAR receptor dependent LTP^23^ this is plausibly an NMDAR-dependent breakthrough of excitation. A specific mechanism that separates females with epilepsy with and without catamenial exacerbation of seizures remains to be found. However, considering the role of NMDARs is an important avenue for research as GABAergic approaches to treating catamenial epilepsy have demonstrated limited efficacy.^9,37^

### Healthy Cohort Computational modelling

The use of DCM may help to unpack the menstrual cycle effect on the underlying neural circuitry changes driving enhanced LTP by detangling to what degree it is co-occurring with increases in NMDAR and AMPAR mediated excitatory parameters or decreases in GABAAR mediated inhibitory parameters.

Feedforward connections between spiny stellate cells (layer IV) and superficial interneurons (layer II/III; (ss→si), as well as superficial interneuron self-gain (si→si) were modulated more by LTP in the perimenstrual phase than the mid-follicular phase. Superficial layer II/III and layer IV have been established as key sites for the induction of LTP in the visual cortex in animal models^38^ Superficial layers II/III contain the highest density of NMDAR in the neocortex, while layer IV is known to be involved in gating mechanism that restrain the activity that can pass to the modifiable synapses in layer II/III.^38^ Therefore, increased intraneuronal input and inhibitory GABAergic self-gain may reflect an overall increase in inhibitory gating, restricting LTP induction through layer IV to superficial layers II/III, which is consistent with the ERP finding of significantly reduced P2 potentiation (as a product of LTP) during the perimenstrual phase, and to a lesser extent, the luteal phase.

During the mid-luteal phase, we found a significant increase in LTP induced modulation of si→si, and excitatory feedforward connectivity from layer IV to layer II/III (ss→si), and excitatory cortico-thalamic projections to relay cells (tp→rl), as well as a significant reduction in the modulation of feedforward connections between layer II/III and layer IV (sp→ss). Modulation of the reciprocal connections between superficial interneurons and spiny stellates, as well as cortico-thalamic projections to relay cells have been implicated previously in mediating LTP changes across the medicated menstrual cycle.^27^ In that study, higher visual LTP was found in the perimenstrual-like phase (the day 1-2 before the hormone-free interval) and was associated with a reduction in these parameters^27^. Here, we found increased modulation of these parameters, except one, during the mid-luteal phase (where there was also reduced P2 potentiation), providing further evidence that these connections are involved in mediating inhibitory gating mechanisms, and are thus inversely related to the magnitude of LTP.

The decrease in inhibitory input to layer IV is unexpected and does not align with the expected enhanced inhibition during the mid-luteal phase^39^ or the findings of Stone, Alshakhouri, Shaw, Muthukumaraswamy and Sumner ^27^. However, this unexpected seemingly contradictory finding could explain why LTP magnitude was only moderately and non-significantly reduced at the ERP level. Secondly, while this is the second study to report the involvement of cortico-thalamic projections to relay cells in mediating LTP changes across the menstrual cycle, this parameter was not found to be modulated by the task itself.^31^ This may suggest that modulation of this parameter is specific to varying the menstrual cycle phase. Supporting this, corticothalamic synapses have high NMDAR/AMPAR density,^40^ which may make them sensitive to the large shifts in oestradiol across the menstrual cycle.

### Epilepsy Cohort Computational modelling

In the epilepsy cohort, LTP-induced modulation of superficial pyramidal cells inhibitory GABAergic self-gain (sp→sp), excitatory output to deep pyramidal cells (sp→dp) and tp→rl were reduced during the perimenstrual phase relative to the mid-follicular phase. Consistent with the control cohort, the modulation of spiny stellate cells feedforward connections to superficial interneurons (ss→si) was increased in the perimenstrual phase as well as in the mid-luteal phase relative to the mid-follicular phase.

In a previous application of the same model, the LTP task induced an increase in sp→sp and a decrease in sp→dp.^31^ Sp→sp represents an inhibitory connection and can therefore be thought of as self-inhibition of superficial pyramidal cells.^31^ In the current study, we found a reduction in the modulation of sp→dp during the perimenstrual phase where P2 potentiation was enhanced. However, we found the enhanced LTP during this phase to be associated with reduced modulation of sp→sp. While this finding seems to contradict previous findings,^31^ conceptually reduced self-inhibition of superficial pyramidal cells appear to align with the measured enhancement of P2 potentiation at the ERP level. Future research ought to explore if this is specific to perimenstrual catamenial epilepsy where GABAergic inhibition is thought to be reduced during the perimenstrual phase.

A reduction in modulation of tp→rl was also found in the epilepsy cohort and co-occurred with enhanced P2 potentiation in the mid-luteal phase. In the control cohort, an increase in the tp→rl parameter co-occurred with reduced P2 potentiation during the mid-luteal phase. Indicating this parameter may be directly related to the magnitude of visual LTP and capture the difference between the control and epilepsy cohorts. Again, this parameter may be of use in future research into perimenstrual catamenial epilepsy and changes in females with epilepsy across the menstrual cycle.

Excitatory feedforward input from spiny stellate cells to superficial interneurons (ss→si) was modulated significantly more during both the perimenstrual phase and the mid-luteal phase compared to the mid-follicular phase in both cohorts also despite opposing ERP results. Of note, the increase in the modulation of this parameter in the healthy cohort was accompanied by increased interneurons inhibitory self-gain which might have resulted in an overall enhanced inhibition and therefore the observed reduction in LTP during the perimenstrual and mid-luteal phases.

Overall, LTP was found to be differentially modulated by the menstrual cycle in females with and without epilepsy and this differential modulation seems to be underlined by differences in the way the brain responds to the same endocrine events. We found no significant differences in hormone absolute levels or relative ratios between the two cohorts, yet the DCM revealed significant differences in the microcircuitry driving the measured LTP. The mid-follicular phase was associated with the most enhanced LTP in the control cohort as expected, but with the least enhanced LTP in the epilepsy cohort. Our findings seem to suggest that while some reduction in inhibition seems to be implicated during the perimenstrual phase, enhanced LTP during the mid-luteal and perimenstrual phases appears to be primarily mediated by upregulation of oestradiol-related excitation. Parameters sp→sp, sp→dp, and tp→rl in particular warrant further use in studies narrowing in on catamenial seizures.

### Limitations

While we were unable to match the numbers in the epilepsy and control cohort, Fig. 4 demonstrated that the finding that visual LTP was weakest at the group level in the mid-follicular phase of females with epilepsy was robust across individuals. It was also consistent with findings using TMS and PPI in epilepsy.^10–13^ The number of participants in this study was too low to fully explore catamenial epilepsy and visual LTP. For example, by exploring potential differences in the model parameters depending on classification. It is also evident that there are challenges with classifying catamenial epilepsy outside of the most severe cases. Recruitment for this study is ongoing with the intention of having sufficient numbers in several years to carry out catamenial epilepsy classification-driven investigations.

## Conclusion

In summary, this study examined menstrual cycle-related changes in visually induced NMDAR-dependent LTP, in females with and without epilepsy. The study demonstrates the perimenstrual phase is a distinct phase, most like the mid-luteal phase. The findings indicate that although females with epilepsy show a markedly different pattern of LTP driven P2 ERP modulation compared to controls over the menstrual cycle, this difference appears to be independent of catamenial seizure pattern. The result suggests that in epilepsy there is an increase in NMDAR driven excitation, driven by oestrogen, that breaks through the seizure protective effects of allopregnanolone of the mid-luteal and perimenstrual phase. Future research should investigate in humans how, or if, this is accompanied by dysfunction to allopregnanolone and GABAAR-δ mediated inhibition or whether oestradiol is the key target.

## Data availability

The data that support the findings of this study are available from the corresponding author, upon reasonable request.

## Supplementary material

Supplementary material is available at *Brain* online.

