## Supplementary Material for "Females with epilepsy show abnormal changes to perimenstrual sensory induced long-term potentiation"

### Methods

#### Hypotheses

(1) The luteal phase would show reduced LTP-induced modulation of the VEP compared to the follicular phase for females with and without epilepsy. Thus demonstrating the dominance of ALLO driven inhibition.
(2) The perimenstrual phase would be associated with less visual LTP than the mid-follicular phase, but greater than the luteal phase in all cohorts. Thus demonstrating sensitivity to ALLO withdrawal reducing GABAergic inhibition and allowing breakthrough of E2 excitation. Exploratory analyses of people with C1 catamenial epilepsy would show they have the greatest visual LTP in the perimenstrual phase.
(3) Computational modelling will show changes in parameters consistent with increased GABAergic inhibition in the mid-luteal phase compared to the mid-follicular phase for all cohorts. There will be a shift to increased NMDA driven excitation and decreased GABAergic inhibition in the perimenstrual phase.

#### Inclusion and exclusion

Participants were excluded if pregnant, taking any hormonal medication that suppresses or interferes with the menstrual cycle, have ever undergone gender-affirming surgical or hormone treatment procedures, or have any current or previous diagnosis of menstrual cycle-related disorders, including symptomatic polycystic ovarian syndrome (PCOS), premenstrual dysphoric disorder, or menstrual migraines. They were also excluded if they have been regularly using any other medication the research team considers may be a problem for the study measures (to be considered on a case-by-case basis and includes regimens like tapering a benzodiazepine around the menstrual cycle). Participants with epilepsy were additionally deemed ineligible if they had a diagnosis of contrast photo-sensitive epilepsy, less than one seizure per month at the time of screening or had damage to their visual cortex confirmed by MRI or determined to be highly likely by a neurologist due to the type of epilepsy or to visual symptoms.

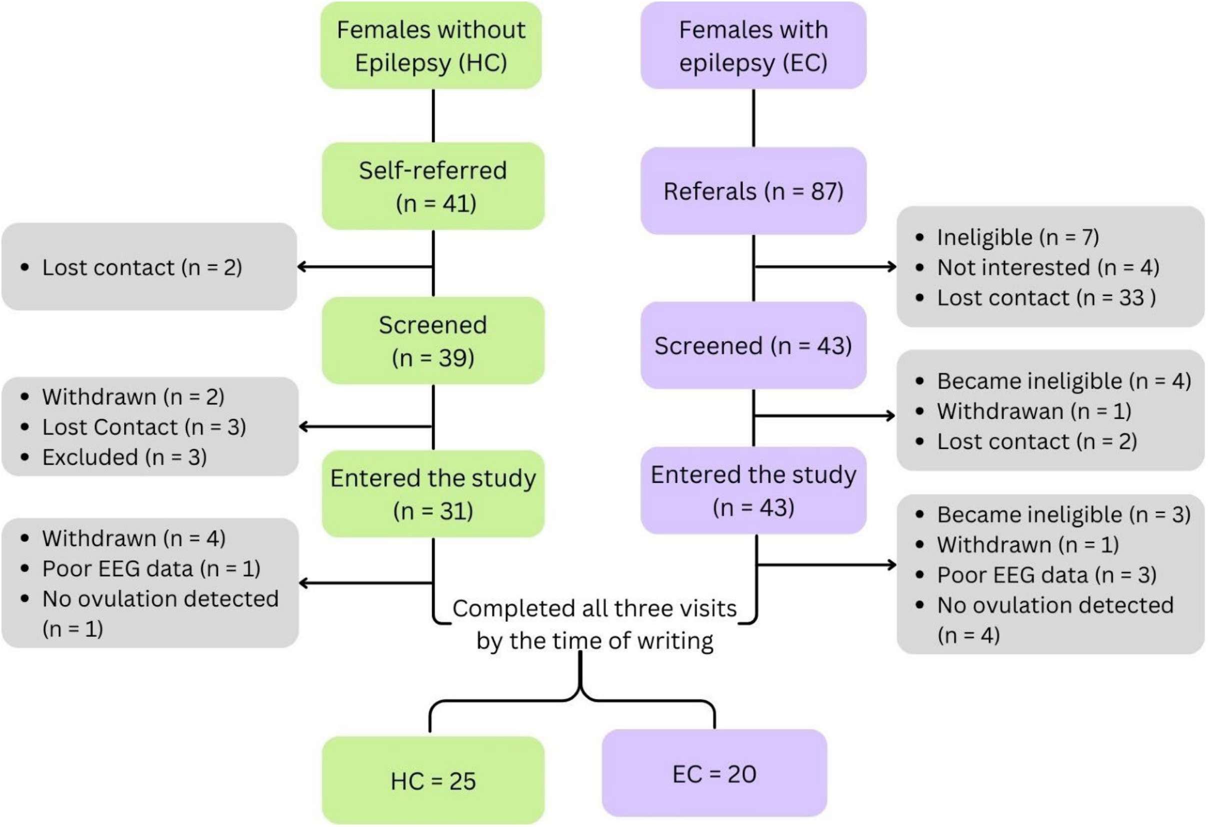


**Figure S1 A CONSORT diagram of sample size from screening to study completion.**

### Blood sampling

Blood samples were collected at each session using 4.5ml PST™ II BD vacutainer® tubes. The PST tubes were refrigerated immediately at 4°C for up to 24 hours before being taken to LabPlus (Auckland Hospital) for quantification of oestradiol and progesterone concentration in plasma by electrochemiluminescence immunoassay. Assays were performed according to Roche Progesterone III (2021) and Oestradiol III (2021) assay guidelines using a Roche Cobas 8000 analyser (e801 module).

### EEG Protocol

EEG data were recorded in Brain Vision Recorder (Brain Products GmbH, Germany) at a sampling rate of 1000 Hz, a resolution of 0.1 µV and a 0.1 Hz low pass filter. Electrodes FCz and FPz were used as online reference and ground, respectively.

Visual stimuli were displayed on an ASUS VG248QE computer monitor with a screen resolution of 1920 x 1080 and 144 Hz refresh rate. Stimulus events were synchronised with EEG through transistor-to-transistor logic (TTL) pulses generated through the parallel port of the display computer. Participants were seated 90cm in front of the display monitor and the distance was checked between recordings.

The stimulus has a spatial frequency of 1 cycle per degree, presented at full contrast on a grey background and subtending 8 degrees of visual angle. Participants were seated 90cm away from the screen and were instructed to passively fixate on a red dot presented in the centre of the screen throughout the task, with the distance confirmed prior to each condition.

In the pre-tetanus condition, horizontal and vertical stimuli were presented 120 times each in random order at low frequency (1 Hz), for a total duration of 4 minutes. The purpose of the pre-tetanus condition was to establish a baseline ERP amplitude for subsequent comparison with post-tetanus conditions. The second condition was a 2-minute photic tetanus involving 1000 presentations of either the horizontal or vertical orientation at high frequency (9Hz) to induce LTP. The orientation of the stimulus was counterbalanced between participants to test for input specificity.

for a total duration of 4 minutes. Each stimulus was presented for 34.8 ms, with interstimulus intervals randomised between five intervals from 896 to 1036 ms, which occurred pseudorandomly but equally often.

The 9 Hz frequency was chosen as it has been shown to reliably induce potentiation of the VEP. The interstimulus interval was either 62.6 or 90.4 ms, occurring at random but equally often. The baseline 1Hz frequency stimulation was repeated as an early-post tetanus condition to record short-term potentiation after a 2-minute break and again after 40 minutes as a late post-tetanus condition to record early LTP. The purpose of the break is to allow retinal after-images from the photic tetanus to dissipate and ensure that any effects measured were not merely reflecting general cortical excitability.

**EEG Pre-processing**

Data were first epoched into 700ms time windows (200ms pre- and 500 ms post-stimulus onset), with the 200 ms prior to stimulus onset used for baselining. Ocular artefacts were removed using a semi-automated process, followed by manual artefact rejection. A modified version of the Fieldtrip semi-automated electro-oculogram artefact (EOG) rejection tool (ft_artifact_zvalue) was applied to frontal electrodes Fp1 and Fp2. Additionally, a second-order 0.3-1 Hz Butterworth bandpass filter was applied with 150 ms of negative trial padding to eliminate only trials in which blinks occurred during the stimulus display.

Manual artifact rejection was then used to eliminate any additional EOG artefacts, muscular artifacts such as clenching, and bad electrodes. During this process, trials rejected by the semi-automated process where reviewed and were restored if the tool had been too restrictive. Remaining electrical artifacts were then removed with a 30 Hz low pass filter. Independent Component Analysis (ICA) via the Fieldtrip function ft_componentanalysis, was then used to identify ECG artefacts, and any persisting EOG, which were then removed manually based on topography and time course using ft_rejectcomponent.

Finally, the Fieldtrip function ft_channelrepair was used to replace bad electrodes via spline-interpolation, which interpolates missing data from the surrounding electrodes, before being converted into SPM12 format. The advantage of using SPM is that it represents spatio-temporal data as a continuous statistical parametric map, allowing for testing large ROIs while still controlling for multiple comparisons. In doing so, it identifies clusters of significance across both time and space, ensuring objective identification of time windows for subsequent analyses.

The pre-processed data were then averaged based on whether they were tetanised or non-tetanised for each session at each of the recording conditions (pre-tetanus, early post-tetanus, and late post-tetanus). The trial averages were then converted into NIfTI images using a time window of 0-250ms and smoothed using a 6 x 6 x 6 x FWHM Gaussian kernel. Based on previous LTP research[1-3] an occipital-parietal region of interest (ROI) was defined, consisting of electrodes P1, P2, P3, P4, P5, P6, P7, P8, Pz, PO3, PO4, PO7, PO8, PO9, PO10, POz, O1, O2 and Oz. In preparation for analysis, individual difference waves for the early (early post-tetanus minus pre-tetanus) and late (late post-tetanus minus pre-tetanus) recordings of each session were then computed using spm_imcalc.

**Table 1.2 Contrasts for the parameter finding analysis.**

| Target test | Direction | Location | SPM contrast |
|---|---|---|---|
| Early potentiation | Negative | Bilateral | -1 -1 0 0 -1 -1 0 0 -1 -1 0 0 |
| Late potentiation | Positive | Central | 0 0 1 1 0 0 1 1 0 0 1 1 |
| Early specificity | Any | Any | 1 -1 0 0 1 -1 0 0 1 -1 0 0 |
| Late specificity | Any | Any | 0 0 1 -1 0 0 1 -1 0 0 1 -1 |

*Note:* FWE-c = Family-wise error corrected. SPM contrasts are conducted according to the following files order: (1) follicular early non-tetanised, (2) follicular early tetanised, (3) follicular late non-tetanised, (4) follicular late tetanised, (5) luteal early non-tetanised, (6) luteal early tetanised, (7) luteal late non-tetanised, (8) luteal late tetanised, (9) perimenstrual early non-tetanised, (10) perimenstrual early tetanised, (11) perimenstrual late non-tetanised, (12) perimenstrual late tetanised.

### Source analysis

Group inverse reconstruction analysis involved using Multiple Sparce Priors method in SPM12. A standard 64-channel actiCAP template of electrode locations with standardised fiducials was used, with SPM's in-built Montreal Neurological Institute (MNI) structural template used in the absence of individual structural MRI scans. A peak voxel was selected from the most significant contrast that encompassed the effects of menstrual cycle on LTP. The selected peak was extracted as virtual local field potential (LFP), with a 5 mm spherical radius around the MNI coordinate. The MNI coordinate was chosen from the most significant peak for each cohort separately and the effects on either hemisphere were assumed to be equal.

***Cohort without epilepsy.*** A time window of 170-250ms was selected for source analysis to capture the menstrual cycle effect on P2 potentiation. Statistical analysis was a 3x3 repeated measure ANOVA ([Phase: mid-follicular vs luteal vs perimenstrual] x [Time: pre-tetanus vs early post-tetanus vs late post-tetanus]). A peak voxel was selected from the most intense F value for a contrast that encompassed both early and late effects of photic tetanus on the mid-follicular and perimenstrual session. This contrast revealed bilateral middle occipital gyrus sources (MNI coordinates left: [-8 -98 -12], right: [26, -98, -8]).

***Cohort with epilepsy.*** Source analysis was run on the same time window as the control cohort (170-250ms) since the cycle effect on P2 potentiation occurred at a similar time (at 183ms for the control cohort and at 199ms for the epilepsy cohort) and using the same statistical design. The only difference was that the peak voxel was selected from the most intense F value for a contrast that encompassed effects of photic tetanus across all three cycle phases given the ERP results that showed that LTP modulation in the mid-follicular phase differed significantly from both the mid-luteal and the perimenstrual phases.

### DCM

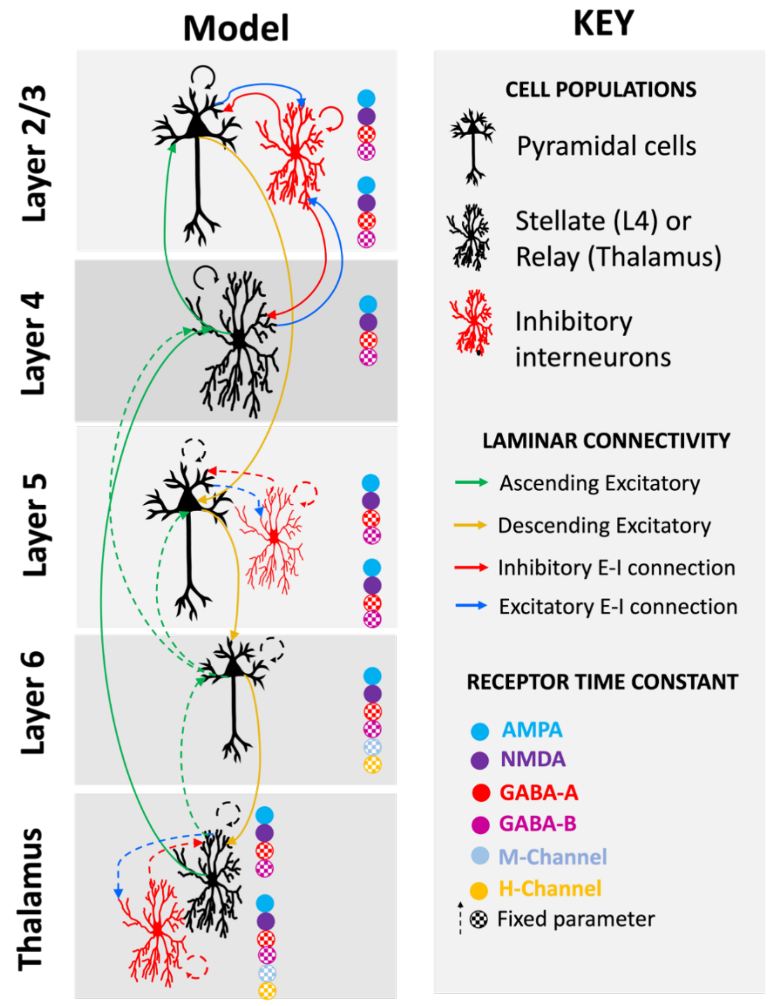


Figure S2 Schematic of Thalamocortical Models (TCM) columnar cell architecture and connectivity. The model consists of 6 cortical neural populations and 2 subcortical (thalamic) populations. Cortical populations include layer 2/3 superficial pyramidal (sp) and superficial interneuron (si) cells, layer 4 spiny stellate cells (ss), layer 5 deep pyramidal (dp) and deep interneuron (di) cells, and layer 6 thalamic projection (tp) cells. Thalamic populations include reticular (rt) and relay cells (rl). Connectivity between cells includes ascending (in green) and descending (in yellow) connections between excitatory cell populations, and inhibitory (in red) and excitatory (in blue) connections between excitatory and inhibitory cell populations. Solid lines indicate connectivity parameters which were allowed to change within the model, and dashed lines indicate fixed parameters. The model also parameterises the decay constants of AMPA, NMDA, GABA-A, GABA-B, M- and H-channels.

The design matrix for the first order PEB consisted of the group averaged effect (represented by a column of 1s), and menstrual cycle effect (represented by a column of 1s for the mid-follicular session and -1s for the perimenstrual session) as well as a random between-subject effects (represented by a column per participant of 1s where their data was). The second order analysis used Bayesian model reduction to re-estimate all the initial parameters that were allowed to vary (often termed the 'full' model (ref)). Bayesian model reduction iteratively searches all possible parameter contributions to the effect of interest, reducing free parameters until only those which meaningfully contribute to the model evidence remain.
